# A Comprehensive Assessment of Somatic Mutation Calling in Cancer Genomes

**DOI:** 10.1101/012997

**Authors:** Tyler S. Alioto, Sophia Derdak, Timothy A. Beck, Paul C. Boutros, Lawrence Bower, Ivo Buchhalter, Matthew D. Eldridge, Nicholas J Harding, Lawrence E. Heisler, Eivind Hovig, David T. W. Jones, Andrew G. Lynch, Sigve Nakken, Paolo Ribeca, Anne-Sophie Sertier, Jared T. Simpson, Paul Spellman, Patrick Tarpey, Laurie Tonon, Daniel Vodák, Takafumi N. Yamaguchi, Sergi Beltran Agullo, Marc Dabad, Robert E. Denroche, Philip Ginsbach, Simon C. Heath, Emanuele Raineri, Charlotte L. Anderson, Benedikt Brors, Ruben Drews, Roland Eils, Akihiro Fujimoto, Francesc Castro Giner, Minghui He, Pablo Hennings-Yeomans, Barbara Hutter, Natalie Jäger, Rolf Kabbe, Cyriac Kandoth, Semin Lee, Louis Létourneau, Singer Ma, Hidewaki Nakagawa, Nagarajan Paramasivam, Anne-Marie Patch, Myron Peto, Matthias Schlesner, Sahil Seth, David Torrents, David A. Wheeler, Liu Xi, John Zhang, Daniela S. Gerhard, Víctor Quesada, Rafael Valdés-Mas, Marta Gut, Thomas J. Hudson, John D. McPherson, Xose S. Puente, Ivo G. Gut

## Abstract

The emergence of next generation DNA sequencing technology is enabling high-resolution cancer genome analysis. Large-scale projects like the International Cancer Genome Consortium (ICGC) are systematically scanning cancer genomes to identify recurrent somatic mutations. Second generation DNA sequencing, however, is still an evolving technology and procedures, both experimental and analytical, are constantly changing. Thus the research community is still defining a set of best practices for cancer genome data analysis, with no single protocol emerging to fulfil this role. Here we describe an extensive benchmark exercise to identify and resolve issues of somatic mutation calling. Whole genome sequence datasets comprising tumor-normal pairs from two different types of cancer, chronic lymphocytic leukaemia and medulloblastoma, were shared within the ICGC and submissions of somatic mutation calls were compared to verified mutations and to each other. Varying strategies to call mutations, incomplete awareness of sources of artefacts, and even lack of agreement on what constitutes an artefact or real mutation manifested in widely varying mutation call rates and somewhat low concordance among submissions. We conclude that somatic mutation calling remains an unsolved problem. However, we have identified many issues that are easy to remedy that are presented here. Our study highlights critical issues that need to be addressed before this valuable technology can be routinely used to inform clinical decision-making.

**Abbreviations and Definitions:**

SSM: Somatic Single-base Mutations or Simple Somatic Mutations, refers to a somatic single base change
SIM: Somatic Insertion/deletion Mutation
CNV: Copy Number Variant
SV: Structural Variant
SNP: Single Nucleotide Polymorphisms, refers to a single base variable position in the germline with a frequency of > 1% in the general population
CLL: Chronic Lymphocytic Leukaemia
MB: Medulloblastoma
ICGC: International Cancer Genome Consortium
BM: Benchmark

aligner = mapper, these terms are used interchangeably

## Introduction

The International Cancer Genome Consortium (ICGC) is characterizing over 25,000 cancer cases from many forms of cancer. Currently there are 74 projects supported by different national and international funding agencies. As innovation and development of sequencing technologies drove prices down and throughput up, projects transitioned from exome to whole genome sequencing (WGS) of tumor and matched germline samples, supplemented by transcript and methylation analyses when possible. These studies have discovered new biology for many different forms of cancer. Interestingly, even though WGS is used for many of these studies, the analyses to date have focused on protein-coding exons that represent only 1-2% of the genome. Data and results produced by ICGC partners are deposited in a common database (https://dcc.icgc.org/). The ICGC has established working groups to empower cross-learning among the various projects and their staff. It became apparent early on that there were marked differences in how teams generated WGS data and analyzed it. In order to understand some of the issues emanating from different sequencing and analysis approaches, the ICGC Verification Group was created and several cross-centre comparisons, called benchmarks (BM), were launched.

Based on cost, capacity and analytical experience it was determined that comprehensive identification of tumor-specific somatic mutations requires WGS with a minimum of 30x sequence coverage of the tumor and normal genomes^11^. Currently, next generation sequencing generates billions of short reads of ~100 bases in length per sample (read length is increasing), which are then typically aligned to a reference human genome. Differences between tumor and normal genomes are identified as Somatic Single-base Mutations (SSM), Somatic Insertion/deletion Mutations (SIM) and structural changes (rearrangements and chromosome segment copy number changes). Conceptually this seems like a simple task, however, cancer genome analysis is challenging and subject to multiple pitfalls. The ICGC Technology Working group’s goal is to explore the various analytical methods and develop a set of best practices to share with the research community. How somatic mutation calling is done is crucial and key to the quality of the results that are produced and deposited in the ICGC database. More importantly, we argue that dissemination of this knowhow beyond the research community and its adoption by health care organizations will be a necessary step toward turning high-resolution cancer genome profiling into a robust clinical tool.

Here we present two consecutive benchmarks of somatic mutation calling that describe a two-year examination of the issues of somatic mutation calling from WGS and the lessons learned. In each benchmark, unaligned sequence reads of a tumor and its corresponding normal genome were made available to members of the ICGC consortium, who returned somatic mutation calls (SSMs, SIMs and structural mutations). The lists of mutations from the ICGC groups were compared to one another and to experimentally verified mutations. All participants were blind to the status of true mutations, as the verification of mutations was carried out *post hoc*. Importantly, the objective was not to identify a winner but rather to identify outstanding issues in somatic mutation analysis and begin to formulate and promote a set of best practices to be adopted more widely by genome researchers. It is also important to stress that this was done using data from biological samples corresponding to two different cancer genomes, in contrast to using simulated data.

## Results

Our benchmark of somatic mutation calling was carried out in two sequential phases, BM1.1 on chronic lymphocytic leukaemia (CLL) and BM1.2 on medulloblastoma (MB). The experience gained from the organisation of BM1.1 was used to improve BM1.2 (see Supplemental Note).

For BM1.1, Illumina sequence corresponding to 40x coverage of a CLL tumor sample and the corresponding normal sample were made available to members of the ICGC (see Online Methods for a detailed description). We received 19 SSM and 16 SIM submissions, of which several were independent submissions by the same group using different versions of their pipelines (*e.g.* CLL. C1 and CLL. C2, CLL. D1 and CLL. D2) or a pipeline used for the BM1.2 (see below) that was run later on the BM1.1 (CLL. F, CLL. L and CLL. N, for example). We also received 10 structural variant submissions.

Verification of mutation calls and generation of a “Gold Set” was carried out after the first 14 submissions. Two independent target capture experiments were carried out (**Online Methods**) using as input all somatic mutations (SSM and SIM) called by at least two submitters and up to 500 private SSM calls per submitter from the set of 14 submissions received by that time (see **SupplementalTable 1**). The target design, capture and sequencing were done at two different sites (University of Oviedo, Spain and OICR, Canada) on an IonTorrent PGM and an Illumina MiSeq, respectively, using the same input DNA that was used for the generation of the WGS data. To establish a Gold Set of mutations for CLL the two Haloplex enrichment datasets were merged with the original Illumina sequencing data. Gold Set mutations were classified (**Table 1**) according to the potential issue it might cause in establishing a correct call: Tier 1 mutations have a mutant allele frequency (AF) >= 10%, Tier 2 mutations have AF>=5%, Tier 3 includes all verified mutations supported by unambiguous alignments, while Tier 4 includes those additional mutations with more complicated or ambiguous local alignments and Tier 5 includes those with unusually high or low read depth. The CLL Gold Set had a total of 1507 bona fide mutation calls across all, with 1143, 1239, 1313, and 1320 SSMs in Tiers 1, 2, 3 and 4, respectively, and 118 and 134 SIMs in Tiers 1 and 4, respectively. This corresponds to a mutational load of almost one verified mutation every two Mbp.

**Table 1.**
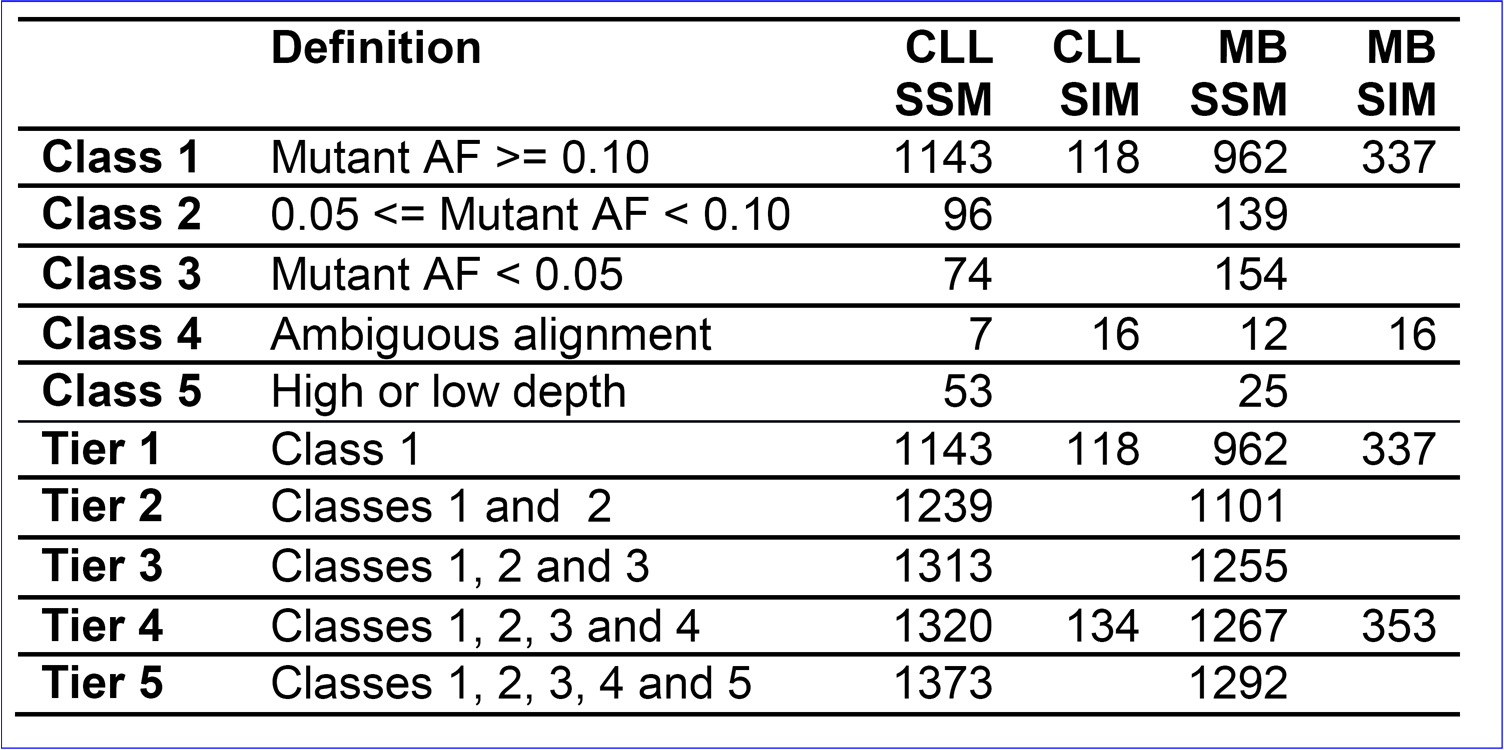
Classification of SSM and SIM Gold Set mutations for both CLL and MB benchmarks.Numbers of curated mutations falling in each class or tier are shown. Successive tiers represent cumulative addition of lower allele frequency mutations, followed by those supported by ambiguous alignments, and finally those with either too low or too high a depth. SIMs were not subjected to such fine classification, with calls only assigned to classes 1 and 4.

During the first phase of the somatic mutation calling benchmark, we initiated a second benchmark on sequencing a medulloblastoma tumor-normal pair (BM2, reference co-submitted manuscript). We realized that while the quality of the reads that had been supplied for BM1.1 was reasonable for the time at which they had been generated, the quality of the sequencing itself had improved substantially for BM2. The ability to call somatic mutations will intimately depend on the quality of the sequences that are used. Consequently we initiated BM1.2 and supplied high quality reads generated by one of the submitters to BM2 to the ICGC somatic mutation calling community for the second part of this benchmark. This time no confidence threshold was prescribed. A similar submission and revision process was set up for BM1.2, but previous experience plus automatic validation and feedback from the submission server reduced the number of validation errors, resulting in more consistent call rates and higher concordance. We received 18 SSM and 16 SIM submissions for BM1.2 that were analysed the same way as the submission for BM.1.1. Four submitters to BM1.1 did not participate in BM1.2 (CLL. R, CLL. S, CLL. T and CLL. U), and there were four new submitters who had not participated in BM1.1 (MB. H, MB. J, MB. M, and MB. Q).

For verification of the medulloblastoma mutation calls (BM1.2), we used a 300x dataset from BM2 to generate a Gold Set (**Supplemental Methods**). In brief, the 300x dataset was analysed using six different somatic mutation calling pipelines. Agreement of at least 3 pipelines was deemed as a positive (a subset of 10% was reviewed manually to confirm this). All calls made by only two or one pipelines were inspected manually and attributed to the tiers outlined above. The MB Gold Set had a total of 1645 bona fide mutation calls across all tiers (**Table 1**), with 962, 1101, 1255, and 1267 SSMs in Tiers 1, 2, 3 and 4, respectively, and 337 and 353 SIMs in Tiers 1 and 4, respectively. The mutational load of this tumor was also close to 0.5 mutations per Mbp.

## Evaluation

Submissions for each benchmark (referred to in the figures as CLL and MB) were compared amongst themselves and to their respective Gold Set (**Figures 1 and 2**). **Figure 1a** and **1c** show the number of SSM calls made per submission that agree with at least one other submission or to the Gold Set. The number of CLL SSMs called ranged from 250 to 18000 in the first 14 submissions prior to correction. CLL. C2, CLL. K and CLL. U contributed a large number of private calls (see **Supplemental Figure 1**) whereas several other submissions had hardly any private calls. **Figure 1b** and **Supplemental Figure 2** show the situation for CLL SIMs, where the number of calls ranged from 18 (CLL. N) to 4152 (CLL. U). It is clear that there is substantially less overlap among SIM call sets. The degree of agreement among MB calls was slightly higher than for CLL, however, still lower than one would have hoped for with 205 SSM calls and only one SIM call that all submitters agreed on (**Figure 1c and 1d and Supplemental Figures 3 and 4**). As in BM1.1 the submitter with the highest number of private calls continued being the highest in BM1.2 while new submissions like MB. H contributed many private calls.

**Figure 1.**
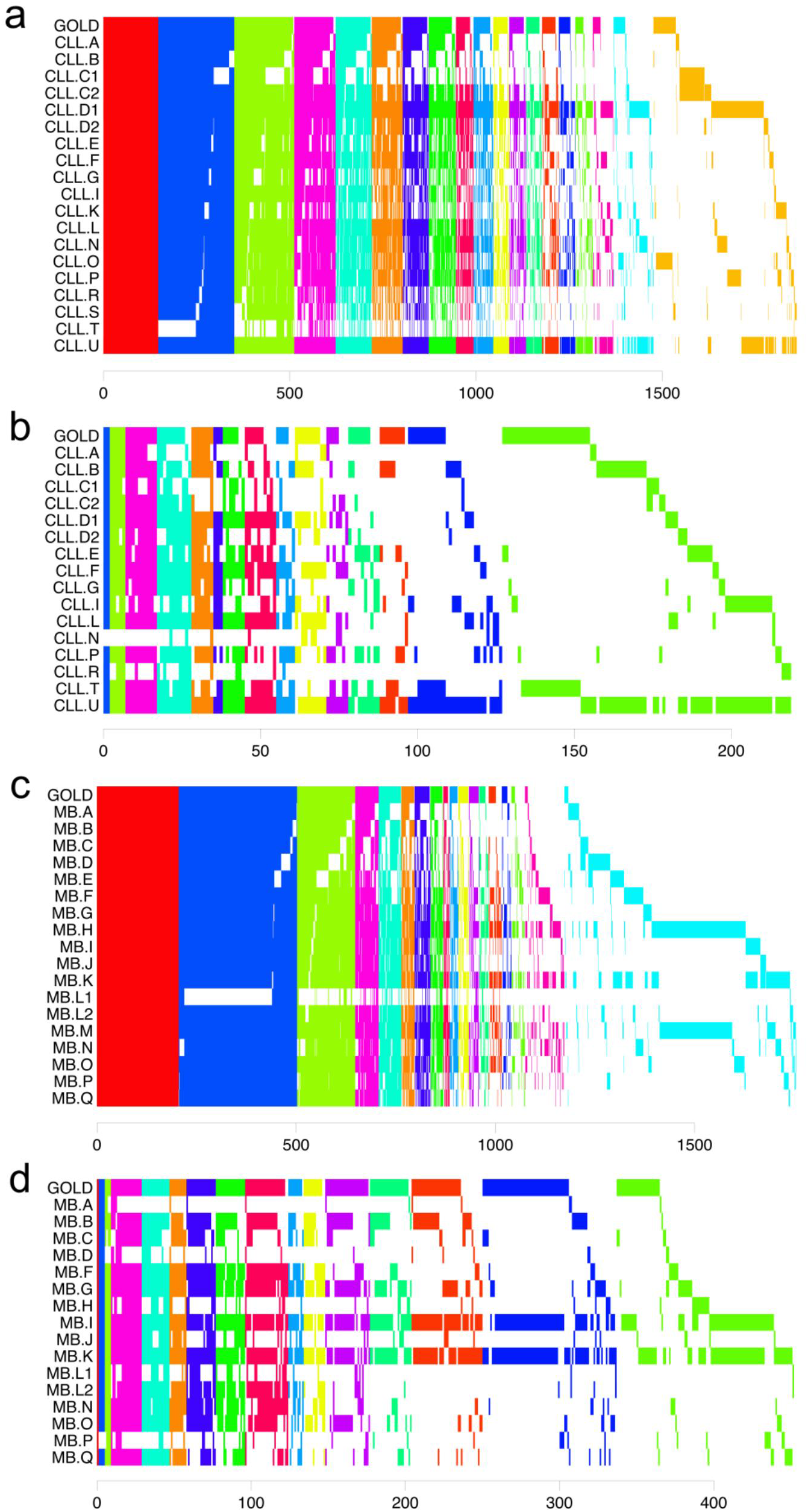
Overlap of somatic mutation calls for each level of concordance. Shared sets of calls arevertically aligned. GOLD indicates the Gold Set. (a) CLL SMM calls shared by at least two call sets. (b) CLL SIM calls shared by at least two call sets. (c) MB SIM calls shared by at least two call sets. (d) MB SIM calls shared by at least two. For all calls see Supplemental Figures 1-4.

In **Figure 2** we show the precision (specificity) of the SSM and SIM calls submitted in function of recall (sensitivity). Best performance is a combination of a high value for precision and a high value for recall (1.0 for recall and 1.0 for precision would be the perfect result). Each letter corresponds to a submission compared to the Gold Set of Tier 1, 2 or 3, with the comparison to Tier 1 (AF>10% mutant allele frequency) having the highest value for recall in the plot and Tier 2 (AF>5%) the middle and Tier 3 (AF>~2%) the lowest. Precision is always calculated against Tier 4, which also includes mutations that are complex or have ambiguous positions. Submissions with a large number of private calls correlate with low precision. For CLL a cluster of submissions achieved recall >0.7 and precision >0.8 for Tier 1. For MB, we observed a similar cluster of well-performing submissions. MB submission with a high number of calls did not necessarily achieve higher recall, however, and the one submission whose precision was the highest (MB. L1) was not significantly more precise than MB. B or MB. Q, which both called over twice as many mutations (**Table 2**). We also looked at the precision and recall of SIMs (**Figure 2b and 2d**). The outcome is rather bleak with no CLL submission achieving a precision or recall greater than 0.8. The submission CLL. T, which displayed high precision but low recall for SSMs, achieved the best balance of precision and recall for SIMs. On inspection of the pipeline used for CLL. T we found that SSMs and SIMs that fell into repetitive elements were eliminated in the filtering step. Some MB SIM submissions achieved precisions greater than 0.9, however their sensitivities were still low.

**Figure 2.**
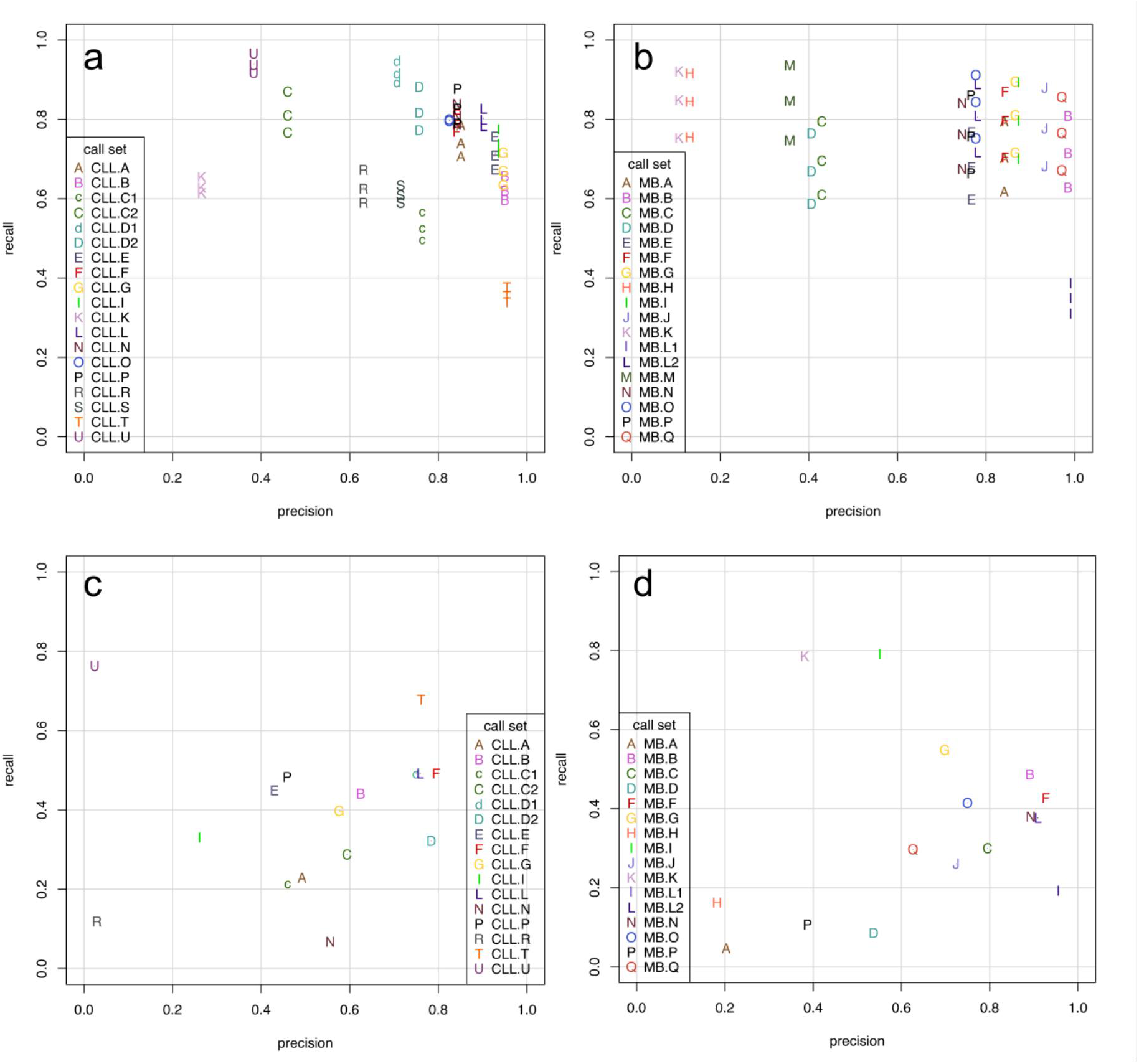
Somatic mutation calling accuracy against Gold Sets. Decreasing sensitivity on Tiers1, 2, and 3 shown as series for each SSM call set, while precision remains the same. a) CLL SSMs. b) MB SSMs. c) CLL SIMs. d) MB SIMs.

**Table 2.** Summary of results. Shown are the evaluation results with respect to the Gold Set as well as our interpretation of the underlying issues affecting performance.

From these results we can make a couple observations. Interestingly the filtering strategy that had been applied by CLL. T that was the most successful for SIM calling and had high precision in BM1.1 is very similar to the one applied by submission MB. L1 for BM1.2. For SSM calls, if CLL. T is removed from BM1.1 and MB. L1 is removed from BM1.2 then the agreement among the rest of the submissions becomes quite good. One of the most specific pipelines but with still decent sensitivity on both benchmarks was CLL. B/MB. B. One of the most sensitive pipelines run on CLL is D1/D2, where the mutation calling method defers alignment to the reference genome until after the differences between the two samples are detected. However, this method performed worse on the MB sample, suggesting that it may be sensitive to the cancer type or mutational signature. The most balanced pipelines for SSMs are those with high F1 scores: CLL. L (0.84), MB. J (0.79) and MB. Q (0.79) (**Table 2**). Interestingly, each of these pipelines combines two different somatic mutation callers, MuTect with Strelka, SGA with FreeBayes, and qSNP with GATK, respectively.

Could those submissions that cluster together in the precision/recall plot (**Figure 2**) be calling the same mutations or have similarities in terms of their pipelines? Using a measure of pair wise overlap, the Jaccard index, we clustered the submissions and display the results as both a heat map and hierarchical clustering in **Figure 3**. Heat maps of just true positive or false positives are provided in **Supplemental Figures 5 to 12**. We find that in general when submissions agree, they tend to agree on true mutations, and when they disagree, these are more likely to be false positives/true negatives. Some notable exceptions can be observed among the false positive CLL SSMs – CLL. E, CLL. G, CLL. I and CLL. B cluster together with a medium amount of overlap, CLL SIMs – CLL. L, CLL. D1, CLL. F and CLL. T cluster, and MB SIMs, where MB. L1 and MB. L2 cluster as do MB. F and MB. N. These correlated false positives may indicate incompleteness of the Gold Sets or just similarities in the pipelines. To address the latter, pipeline features and parameters were converted into discrete values and used to cluster pipelines based on shared features. The clusters found (**Supplemental Figures 13 and 14**) did not mirror the clusters based on the Jaccard index, suggesting that perhaps only a single parameter or filter, not the similarity of the entire pipeline, may be responsible. To focus more on similarities of true positive calls, we performed a correspondence analysis, the results of which are shown in **Figure 4**.

Could those submissions that cluster together in the precision/recall plot (**Figure 2**) be calling the same mutations or have similarities in terms of their pipelines? Using a measure of pair wise overlap, the Jaccard index, we clustered the submissions and display the results as both a heat map and hierarchical clustering in **Figure 3**. Heat maps of just true positive or false positives are provided in **Supplemental Figures 5 to 12**. We find that in general when submissions agree, they tend to agree on true mutations, and when they disagree, these are more likely to be false positives/true negatives. Some notable exceptions can be observed among the false positive CLL SSMs – CLL. E, CLL. G, CLL. I and CLL. B cluster together with a medium amount of overlap, CLL SIMs – CLL. L, CLL. D1, CLL. F and CLL. T cluster, and MB SIMs, where MB. L1 and MB. L2 cluster as do MB. F and MB. N. These correlated false positives may indicate incompleteness of the Gold Sets or just similarities in the pipelines. To address the latter, pipeline features and parameters were converted into discrete values and used to cluster pipelines based on shared features. The clusters found (**Supplemental Figures 13 and 14**) did not mirror the clusters based on the Jaccard index, suggesting that perhaps only a single parameter or filter, not the similarity of the entire pipeline, may be responsible. To focus more on similarities of true positive calls, we performed a correspondence analysis, the results of which are shown in **Figure 4**.

Could those submissions that cluster together in the precision/recall plot (**Figure 2**) be calling the same mutations or have similarities in terms of their pipelines? Using a measure of pair wise overlap, the Jaccard index, we clustered the submissions and display the results as both a heat map and hierarchical clustering in **Figure 3**. Heat maps of just true positive or false positives are provided in **Supplemental Figures 5 to 12**. We find that in general when submissions agree, they tend to agree on true mutations, and when they disagree, these are more likely to be false positives/true negatives. Some notable exceptions can be observed among the false positive CLL SSMs – CLL. E, CLL. G, CLL. I and CLL. B cluster together with a medium amount of overlap, CLL SIMs – CLL. L, CLL. D1, CLL. F and CLL. T cluster, and MB SIMs, where MB. L1 and MB. L2 cluster as do MB. F and MB. N. These correlated false positives may indicate incompleteness of the Gold Sets or just similarities in the pipelines. To address the latter, pipeline features and parameters were converted into discrete values and used to cluster pipelines based on shared features. The clusters found (**Supplemental Figures 13 and 14**) did not mirror the clusters based on the Jaccard index, suggesting that perhaps only a single parameter or filter, not the similarity of the entire pipeline, may be responsible. To focus more on similarities of true positive calls, we performed a correspondence analysis, the results of which are shown in **Figure 4**.

**Figure 3.**
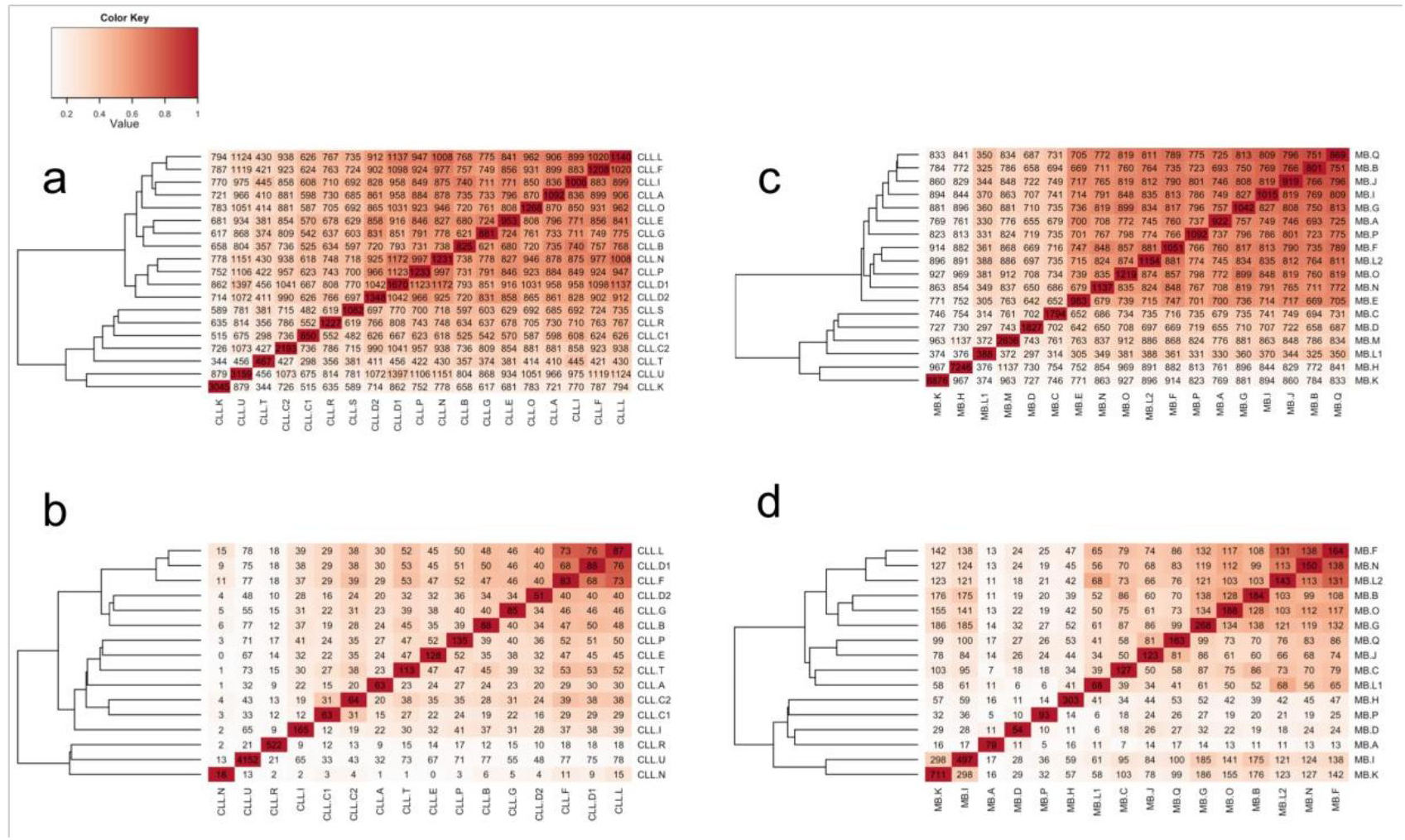
Jaccard index. Heat maps and dendrograms reflect the value of the Jaccard index(pairwise ratio of intersection/union). Values in each cell indicate only the number of somatic mutations shared by each pair of submissions. a) CLL SSMs. b) CLL SIMs. c) MB SSMs. d) MB SIMs.

**Figure 4.**
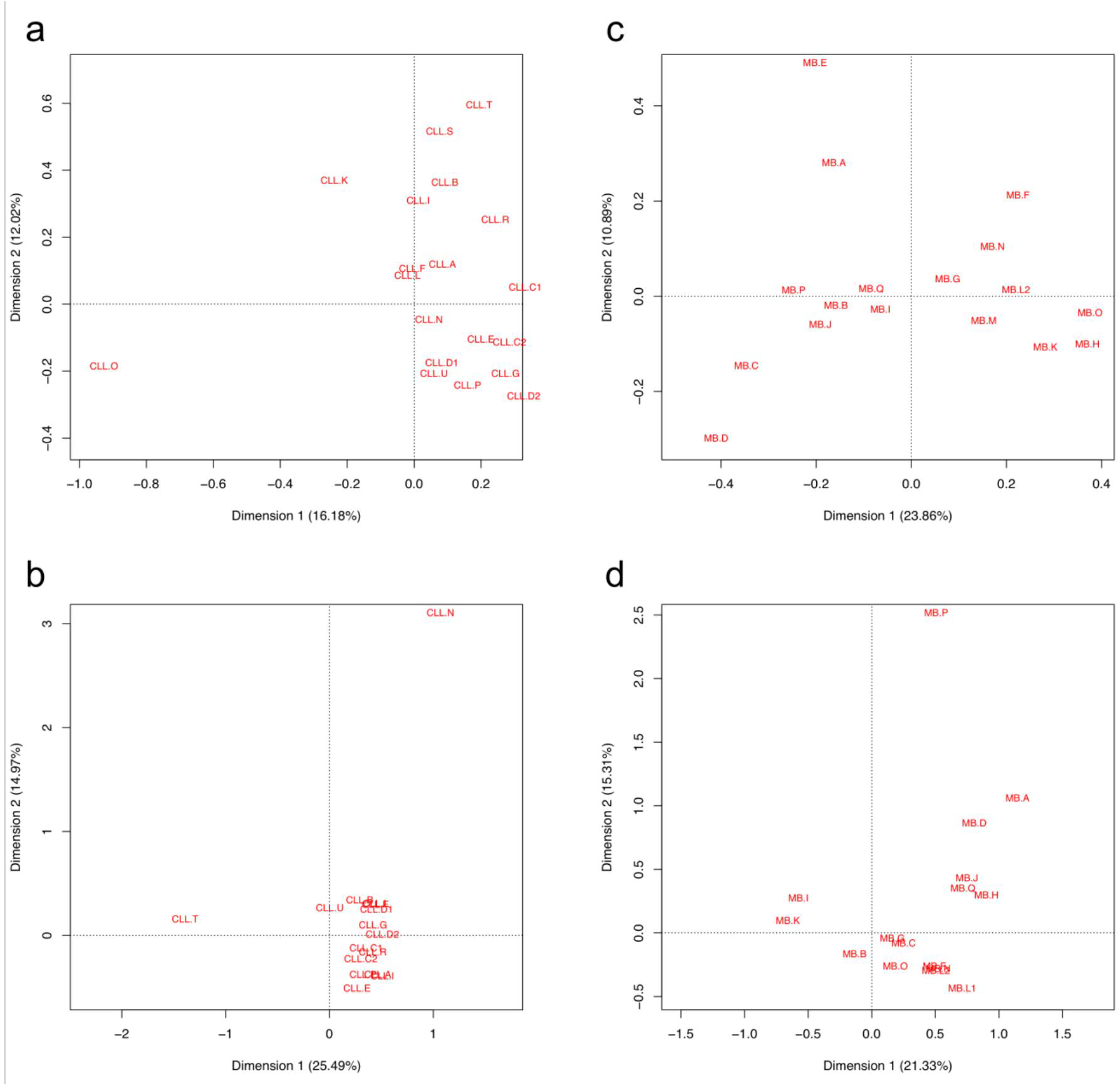
Correspondence analysis of true positive (Gold Set) calls. a) CLL SSMs. b) CLL SIMs. c) MB SSMs. The outlier MB. L1 not shown. d) MB SIMs.

What features correlate with better or worse performance? We know that genomic context plays a role, given that the reads used were relatively short 100 or 150bp reads. So-called “rainfall” plots are a good way at looking at density of features along a genome. By plotting the distance from the previous mutation (**Figure 5 and Supplemental Figures 15 and 16**), we can observe clustering or hotspots. SSM and SIM calls are coloured as true positives, false positives and false negatives. Given that we are looking at only a single cancer case at a time, we do not expect mutational hotspots. However, we detected quite different patterns with submissions like CLL. L that did not show major weaknesses across the genome (**Figure 5a**), while CLL. C2, for example, made many false positive mutation calls in centromeric regions (**Figure 5b**). This latter pattern may arise if alignment of reads is done to the GRCH37 reference without the d5 decoy sequence and no ultra-high-signal blacklist is used. CLL. K dramatically overcalled, with seemingly even coverage across the genome, however in the MB. K rainfall plot (**Supplemental Figure 16**) higher ploidy chromosomes (chr17) display a greater density of calls and lower ploidy chromosomes (8, X and Y) demonstrate a lower density of calls (**Figure 5c**). CLL. R overcalled in regions of chromosomes 11, 13, 14, 15 and 21 while the calls otherwise resembled the pattern of CLL. L (**Figure 5d**). In this case, it would appear that known structural and copy number variation present in this sample affected the local false-positive rates.

**Figure 5.**
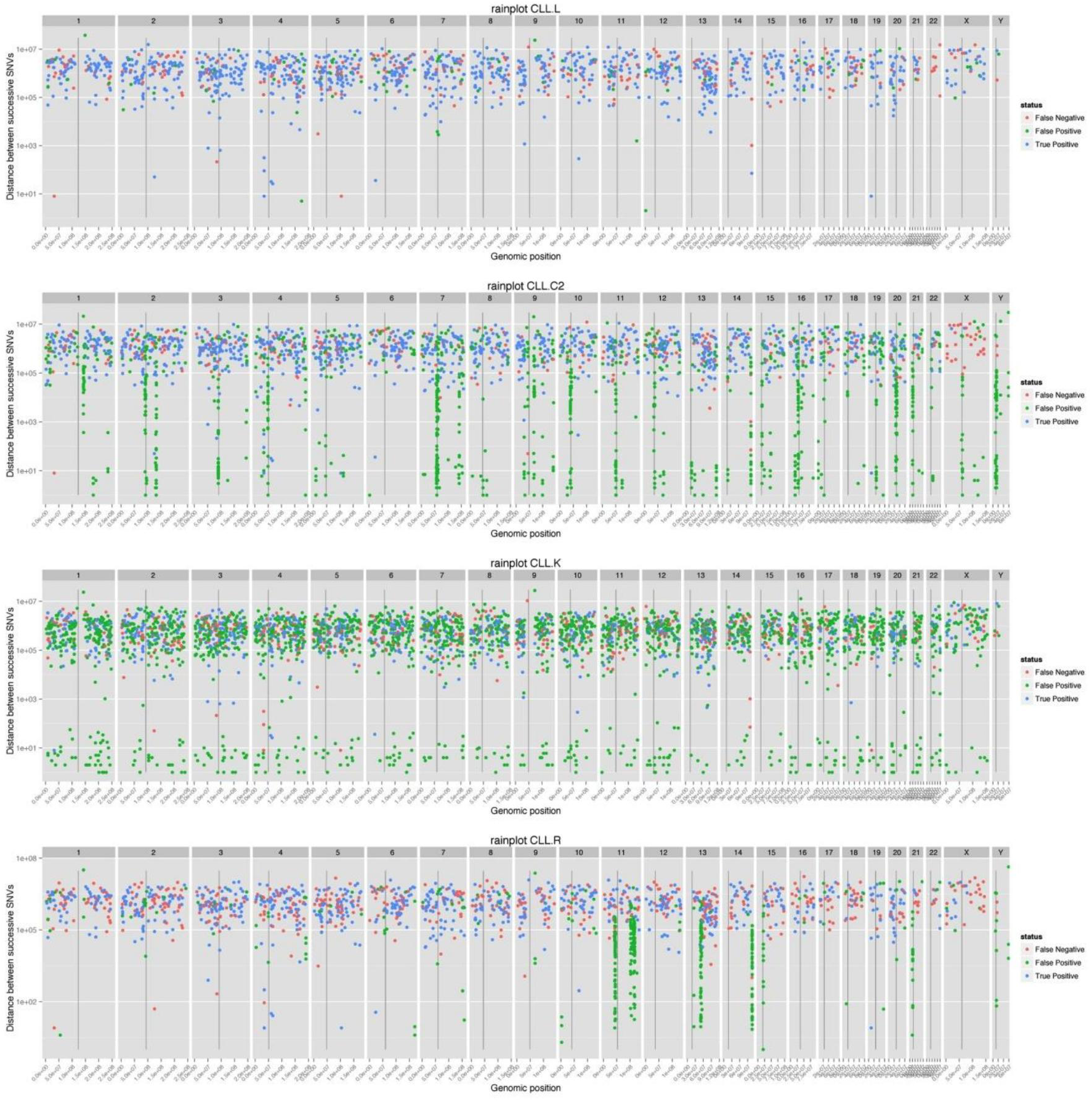
Rainfall plot showing distribution of called mutations on the genome. The distancebetween mutations is plotted in log scale (y axis) versus the genomic position on the x-axis. True positives (blue), false positives (green) and false negatives (red). Four CLL submissions representative of distinct patterns are shown, from top to bottom: CLL. L has low positional bias, CLL. C2 has clusters of false positives near centromeres and false negatives on the X chromosome, CLL. K has a high false positive rate with short distance clustering of mutations, and CLL. R has a concentration of called mutations on chromosomes 11, 13, 14, 15 and 21. The full set of rainfall plots for CLL and MB is in the Supplemental Material.

In another analysis we tried to determine genomic features such as repeat features as well as some key sample/sequence/alignment features that created problems for callers. Membership (Boolean flags) or scores (for continuous characteristics) were tabulated for CLL and MB SSMs and SIMs (**Supplemental Tables 2–5**). The frequencies or mean scores, respectively, were computed for three subsets of each submission (true positives, false positives, or false negatives) and for the Gold Sets. In order to highlight problematic features for each submission, the differences with respect to the Gold Set were computed and multiplied by the false positive or false negative rate, accordingly. The problematic features of false positives in the MB SSM dataset, for which the verification sequence was more uniformly distributed over all candidate mutation loci than that of CLL, are shown as a heat map (**Figure 6**). Nearly all sets of false positives are enriched in low frequency mutations, and thus harder to discriminate from background noise (also reflected by the “sameAF” metric), some call sets (MB. K, H, C, D and M) have more trouble with this category of false positives. MB. H seems to also have a problem with segmental duplications and multi-mappable regions, and MB. D with duplications only. MB. K and MB. M (to a lesser extent) are enriched in SSMs falling in tandem repeats, simple repeats and homopolymers. MB. C has a problem with false positives falling in blacklisted regions, specifically centromeric and simple repeats. Interestingly, the three submissions with fewer false positives immediately adjacent to tandem repeats than in the Gold Set (MB. H, MB. C and MB. D) do not use BWA for the primary alignment step – instead, the mappers Novoalign or GEM are used, or the detection method is not based on mapping (SMUFIN). The corresponding heat maps for MB SSM false negatives, MB SIMs, CLL SSMs, and CLL SIMs are shown in **Supplemental Figures 17 to 23**.

**Figure 6.**
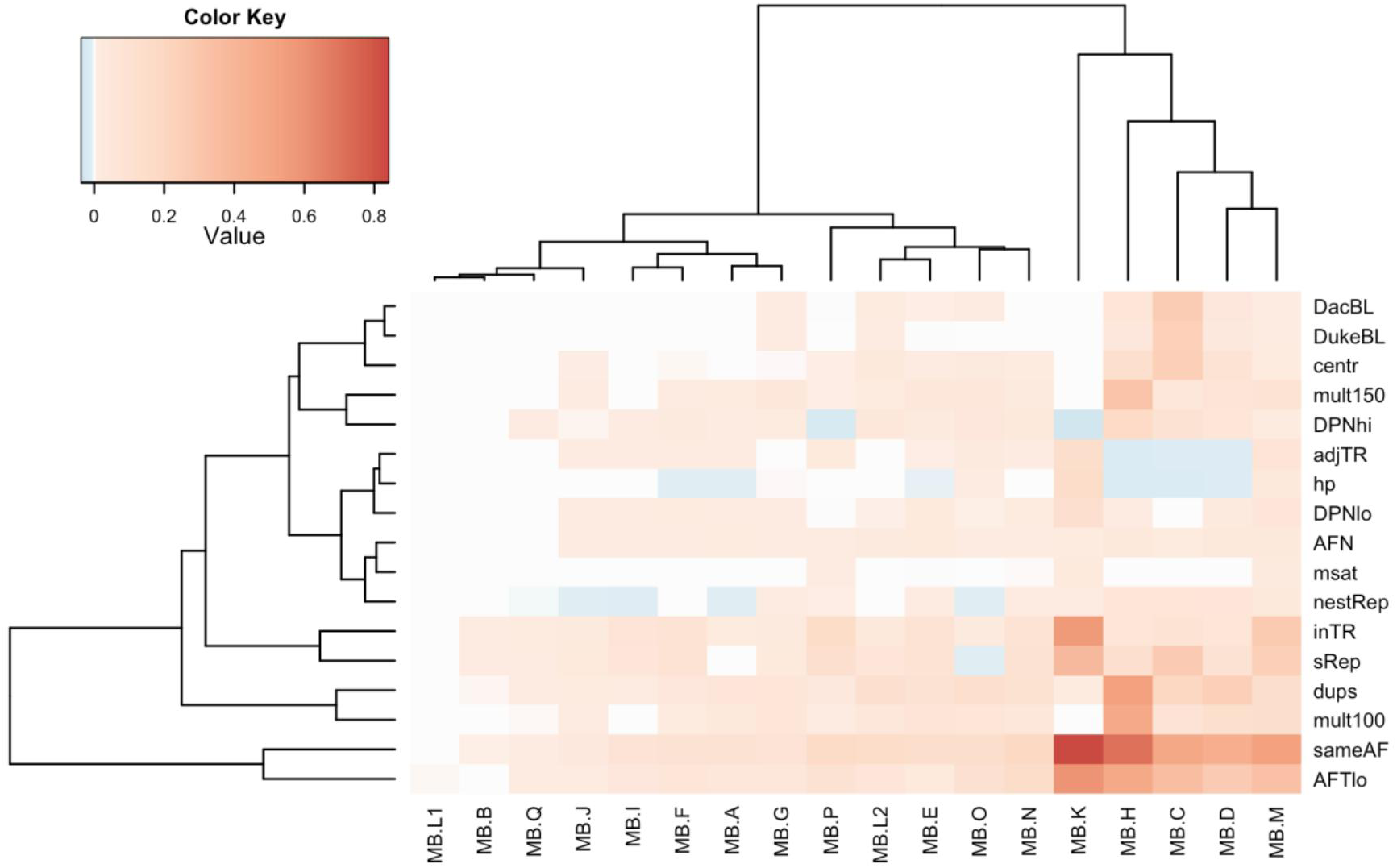
Enrichment or depletion of genomic and alignment features in false positive calls for each MB SSM submission. For each feature, the difference in frequency with respect to the Gold Set is multiplied by the false positive rate. Blue indicates values less than zero and thus the proportion of variants or their score on that feature is lower in the false positive set with respect to the true variants. Reddish colors correspond to a higher proportion of variants or higher scores for the feature in false positive calls versus the Gold Set. Both features and submissions are clustered hierarchically. The features shown here include sameAF (the probability that the allele frequency in the tumor sample is not higher than that in the normal samples, derived from the snape-cmp-counts score), DacBL (in ENCODE DAC mappability blacklist region), DukeBL (in Encode Duke Mappability blacklist region), centr (in centromere or centromeric repeat), mult100 (1–mappability of 100mers with 1% mismatch), map150 (1–mappability of 150mers with 1% mismatch), DPNhi (high depth in normal), DPNlo (low depth in normal), dups (in high-identity segmental duplication), nestRep (in nested repeat), sRep (in simple repeat), inTR (in tandem repeat), adjTR (immediately adjacent to tandem repeat), msat (in microsatellite), hp (in or next to homopolymer of length > 6), AFN (mutant allele frequency in normal) and AFTlo (mutant allele frequency in tumor < 10%). Corresponding plots for SIM, false negative calls and CLL results are in the Supplemental Material.

In addition a correspondence analysis was carried out which confirms the above findings for MB SSM false positives (**Supplemental Figure 24**). MB. C clusters with EncMap and dukeMap, suggesting that MB. C false positives occur in some blacklisted regions. MB. H (and MB. G and MB. O to lesser extent) false positives are associated with segmental duplications. MB. K false positives are associated with tandem repeats (and microsatellites and simple repeats). MB. M and MB. F false positives associate with low depth in the tumor, and MB. L2 and MB. D false positives with high mutant allele frequency. Indeed, there appears (in the rainfall plots, **Figure 5 and supplemental Figures 15 and 16**) to be some overcalling on chromosomes 8 and Y (where allele frequencies are expected to approximate 100%) in MB. D.

The differences between sets of mutations submitted by the participating groups (**Figures 1 and 2**) raised questions about the impact of individual pipeline components on the results. The extent of observed pipeline customization (**Supplemental Note and Supplemental File 1**) did not allow for exhaustive testing of all potentially important analysis steps (even less so with various the parameter settings), but three pipeline components were selected for a closer inspection due to their expected high impact: mapper, reference genome build and mutation caller. Four mappers (Novoalign2, BWA, BWA-mem and GEM), two SSM callers (MuTect and Strelka) and three versions of the human reference genome (b37, b37+decoy and “hg19r” – a reduced version of hg19, with its unplaced contigs and haplotype chromosomes removed) were selected for testing, based on their usage by the benchmarking groups (see **Supplemental Note** for software versions and settings). In order to limit the effect of non-tested software on the produced mutation sets, a simple SSM-calling pipeline was established.

First we compared the effect of the mapper with each of the SSM callers. With a single SSM caller employed, a considerable fraction of unfiltered SSM calls for a given mapper (0.22 to 0.69, depending on the mapper-caller combination), were not reproducible by that caller with any other mapper (**Supplemental Figure 25**). When compared to the Gold Set (Tier 3 SSMs), calls supported by a single mapper were almost exclusively false positives (FP) (precision < 0.02). On the other hand, a large majority of calls supported by all four mappers were true positives (with precision ranging from 0.87 for MuTect to 0.99 for Strelka).

Similar trends could be observed when SSM callers were compared while holding the mapper constant (**Supplemental Table 6**). A sizable fraction (0.22 to 0.87, depending on the mapper) of unfiltered SSM calls for any given mapper-caller combination was not reproducible by the other caller on the same alignment file. When compared to the Gold Set, calls supported by a single caller appeared to be mostly false positives, with precision ranging from 0.01 to 0.05. Calls supported by both callers proved to be mostly correct (with precision between 0.89 and 0.93; **Supplemental Table 7**).

From these numbers we conclude that:

1. Each tested mapper and caller has its own specific biases, which – unless eliminated – might lead to a significant number of false positive calls;
2. Mappers and callers should perhaps not be considered as completely independent components, as certain combinations show much higher compatibility than others (e.g. with Novoalign alignments as input, MuTect produces an SSM call set with the lowest overall precision, while the SSM call set produced by Strelka on the same alignment file has the highest overall precision; table PCT1);
3. Some callers might by default be geared for sensitivity, others for precision; as previously reported, use of a consensus of multiple callers might be a suitable way of elevating precision without making substantial sensitivity sacrifices.

The results of reference genome choice and a detailed examination of the alignment characteristics of the different aligners are presented in the **Supplemental Note**.

## Discussion

This benchmark study shows that, contrary to common perception, identifying somatic mutations, be they SSMs or SIMs, is still a major challenge. This entire benchmark was split into two parts, a first part BM1 on CLL that provided reads of average quality for the time they were produced and a verification strategy of identified mutations with an orthogonal sequencing approach. Several observations were made: 1) Quality of sequencing at the time was still improving dramatically and needed to be accounted for in the mutation calling procedures, 2) we lacked a consensus on boundary conditions, such as nomenclature of somatic mutations, thresholds, the reference to be used and how a result should be reported, 3) the application of an orthogonal method for verification allowed the assessment of false positive reports, however, false negative calls cannot be assessed exhaustively, and 4) the mutation call rates between submission ranged from 467 to 3159 for 1373 verified SSMs and 18 to 4152 for verified 134 SIMs of the Gold Set, respectively. The boundary issues were remedied for BM1.2 on MB. High quality reads were made available that covered the genome evenly and the instructions for reporting were given in even more details. Still the discrepancy between different analysts was quite big with mutation call rates ranging from 801 to 8876 for 1292 verified SSMs and 54 to 711 for 353 verified SIMs of the Gold Set, respectively. In contrast to BM1.1 the Gold Set was established using 300x sequence coverage, which allowed determining sweet spots of pipelines and assessing false negative rates as well as false positive rates. Providing exactly the same sequencing reads from tumor/normal pairs resulted in substantial discrepancies in somatic mutation call rates and the calls themselves in the hands of different analysts. Next we identified the Achilles’ heels of pipelines for failing to identify or overcalling mutations in function of genomic features. We found that dominating features for difficulties both for false positives and false negatives calls for SSMs were low coverage in the tumor, and aberrant coverage of tumor and normal, while for SIM calling repetitive features such as inTRs and sRep tend to dominate both in CLL and MB. We also detected distinct clustering of different analysts. Even though analysts used similar combinations of software in their pipelines, very few similarities were detected in their calls. The combination of software is not as critical as how each piece of software is applied and what settings are applied. A slight correlation of true positive SSM calls of pipelines using MuTect and Strelka can be seen (**Figure 4**). Data analysis pipelines are constructed by integrating software from diverse sources. In many instances the approaches used in the different software and the assumptions made therein are not evident to the user and many parts are black boxes. In order to account for unknowns, pipeline developers resort to calibrating their pipelines against known results, for example from a genotyping experiment done on similar samples, and by using combinations of tools for the same process step, assuming that a result shared by two different approaches has a higher likelihood to be correct. Of note is that many of the pipelines apply multiple tools for the same process step in their pipeline and then use intersects to weight calls. This practice, together with good use of blacklists, results in best outputs. No best tools emerged from our benchmark and it is also clear that no strict best threshold settings exist. However, it is clear that how analysts use their pipeline is critical for the quality of their output. Over the course of the benchmark different analysts were able to improve their calls towards the Gold Sets. So, having a benchmarking dataset is of great importance. The sequencing reads and Gold Sets of BM1.2 are available to the research community through the ICGC DACO to benchmark their own pipelines.

In conclusion, we have conducted a thorough benchmark of mutation calling in cancer genomes. The two tumor types we selected are particular in that they present a comparatively low number of SSMs and SIMs and no structural rearrangements, and the tumor samples we chose had very high tumor cellularity, in the case of the CLL tumor due to cell sorting. The MB tumor added a degree of difficulty to the analysis in that this tumor presented a large degree of tetraploidy combined with other changes of ploidy in chromosomes 1, 4, 8, 15 and 17. As general advice for somatic mutation analysis in cancer genomes we would suggest to start with gaining a basic understanding of nature of the tumor and tumor-type that is being analysed. This allows adapting the downstream analysis to extract the best possible result. Concretely this refers to the order in which analysis should be done, starting with high level characterisation of the large-scale tumor genome features followed by the fine grain analysis of SSMs and SIMs. Also iterative procedures should be considered. Importantly, it should be noted that somatic mutation analysis needs the comparison of the tumor with the normal sample and that a normal sample also needs to be well selected.

Further benchmarks are required to resolve even more difficult mutational features that tumor samples and genomes can present, such as low sample purity (contamination of tumor cells by normal cells and also the normal contaminated by tumor cells or viral components), subclonality, structural rearrangements and karyotype aberrations. Good tools to determine these features prior to the detection of SSMs and SIMs are needed. Resolving issues such as the one that we have elaborated in this benchmark is critical to take cancer sequencing adoptable for the clinic.

## Acknowledgements

Financial support was provided by the consortium projects READNA under grant agreement FP7 Health-F4-2008-201418, ESGI under grant agreement 262055, GEUVADIS under grant agreement 261123 of the European Commission Framework Programme 7, ICGC-CLL through the Spanish Ministry of Science and Innovation (MICINN), the Instituto de Salud Carlos III (ISCIII) and the Generalitat de Catalunya. SD is supported by the Parc Científic de Barcelona through the Torres Quevedo subprogram (MICINN) under grant agreement PTQ-12-05391. The Ontario Institute for Cancer Research has received generous support from the Ontario Ministry of Research and Innovation. Additional financial support was provided by Prostate Cancer Canada, Genome Canada and the Canada Foundation for Innovation. The Synergie Lyon Cancer platform has received support from the French National Institute of Cancer (INCa) and from the ABS4NGS ANR project (ANR-11-BINF-0001-06). The ICGC RIKEN study was supported partially by RIKEN President’s Fund 2011, and the supercomputing resource for the RIKEN study was provided by the Human Genome Center, University of Tokyo. EH is supported by the Research Council of Norway under grant agreements 221580 and 218241 and by the Norwegian Cancer Society under grant agreement 71220-PR-2006-0433. We want to thank Elias Campo for giving us access to the data of the CLL sample for BM1.1 as well as DNA for verification.

### Online Materials and Methods

#### Benchmark 1.1 Sequencing Data – Chronic Lymphocytic Leukaemia

##### Whole Genome Sequencing

Whole genome sequencing reads for BM1.1 were produced using DNA from a patient suffering from CLL who had given informed consent for sample collection and analysis. Tumor samples were from before treatment and tumor cells were separated from non-tumor cells by immunomagnetic depletion of T cells, natural killer cells, monocytes and granulocytes (Puente et al., Nature 2011). Tumor cell purity was >98% as assessed by flow cytometry. Normal blood cells from the same patient were used for the normal sample that contained less than 0.05% tumor cell contamination as assessed by flow cytometry. The tumor sample was sorted using FACS CD. Two different libraries each for the tumor and the corresponding normal DNA sample using Illumina TruSeq™ DNA Sample Preparation procedures with slight modifications. While one library was prepared following the standard protocol with 10 cycles of PCR, the second library was heated to 72˚C before adapter ligation and cooled down suddenly to 4˚C. This resulted in a biased proportion of high GC content reads and counterbalances some of Illumina’s sample preparation methods’ GC-bias (improved coverage of elevated GC-content regions of the genome. The normal and GC enriched libraries were sequenced on Illumina GAIIx (2×150 bp) and Illumina HiSeq2000 (2×100 bp) instruments. The same amount of data (~40x coverage) was produced for tumor and corresponding normal sample with the proportions of the two library types–600 million reads for the standard and 200 million reads for the GC enriched library. Reads in FASTQ format were generated using the RTA software provided by Illumina.

*Verification by target capture and orthogonal sequencing technologies* 4080 SSMs and 883 SIMs were selected for Haloplex target capture. Haloplex target design and capture were performed independently by two different groups, using different sequencing platforms: MiSeq and IonTorrent.

For verification with MiSeq, Haloplex custom reagent was designed for the Illumina HiSeq platform through the Agilent SureDesign portal (https://earray.chem.agilent.com/suredesign). The input contained 4,674 target segments encompassing 192,069bp. Design parameters included an optimal amplicon length of <150bp and each selected variant was padded by 25 bases both 5’ and 3’ to be contained within the assayed segment. Target enrichment was designed from human reference genome hg19 (GRC37, February 2009). The designed Haloplex custom reagent contained 19,392 amplicons encompassing 170,430bp covering 88.73% of the input target. Samples were processed using 200ng of input DNA following the manufacturer provided Haloplex protocol. Each sample was sequenced using the Illumina MiSeq platform with 2 x 150 base reads. 3.3 and 5.0 million reads were obtained covering the Haloplex products for the reference and tumor samples, respectively. Greater than 80% of the original 4,674 input targets were sequenced at over 20x coverage.

For verification of somatic mutations using IonTorrent, a HaloPlex custom capture was designed to enrich a total of 5179 mutations (4492 substitutions and 687 small indels) with a target region of 100 bp centered on the mutation position using the SureDesign software (Agilent). Sequencing was performed using two runs of the IonTorrent 418 chip, resulting in a total of more than 5 and 8 million reads for the tumor and normal samples, respectively.

All 40x reads (GAIIx and HiSeq2000 reads that were provided to the participants), MiSeq and IonTorrent verification reads were aligned with GEM (gem-mapper) and converted to BAM format. Alignments were filtered to retain only primary alignments with mapping quality >= 20. Duplicates were removed with Picard, indels realigned at 1000 genomes indel target locations, and indels were left-aligned using GATK. The pileups at SSM positions were extracted using samtools mpileup with base quality threshold >= 13. Read depth and base counts were extracted using a custom script. Mutant allele and normal counts were compared using in-house software snape-cmp-counts^13^ which compares alternate and reference allele counts in tumor and normal, and then scored according to the probability that they are derived from different beta distributions. Mappabilities with 0, 1 and 2% mismatches were computed for the reference genome (h37d5). The average mappabilities in 100bp windows preceding each candidate mutation were stored as tracks for visualization in IGV. In addition, the segmental duplication annotation from the UCSC browser was loaded into IGV. Mutations were then classified as follows. Mutations with sufficient depth (>=20) and a snape score >=0.98, average mappability of one, and no overlap with segmental duplications were automatically classified in the Gold Set according to their mutant allele frequency (class 1: MAF>=0.1, class 2: 0.1>MAF>=0.05 or class 3: MAF<0.05). All other candidates with snape score >0.9 were reviewed visually in IGV. Mutations with ambiguous alignments were assigned to class 4. Low depth but otherwise plausible mutations were assigned to class 5. Somatic mutation Gold Set tiers were compiled by cumulative addition of classes so that Tier 1 only includes class 1, while Tier 2 includes class 1 and class 2, Tier 3 includes classes 1, 2 and 3, etc. All other candidate mutations were rejected and assigned to class 0 (**Table 1**).

### Benchmark 1.2 Sequencing Data – Medulloblastoma

#### Whole Genome Sequencing

The sequencing reads for BM1.2 were produced using a no-PCR library preparation procedure that was adapted from the KAPA Library Preparation Kit protocol used together with Illumina TruSeq adaptors and omitting PCR amplification, each for the tumor and the corresponding normal DNA sample. For each sample two libraries were prepared with smaller (roughly 300 bases) and larger (roughly 450 bases) insert size. Sizing was done by agarose gel separation and excision of corresponding size bands. The four libraries were sequenced to 40x using Illumina HiSeq2000 (2x100 bp) and 4x using Illumina MiSeq (2x250 bp). Reads in fastq format were generated using the RTA software provided by Illumina.

#### Verification by 300x Coverage

For verification of the submissions of BM1.2 all reads produced for BM2.0 were combined and analysed to generate a curated set of results (Gold Set). The combined sequences gave roughly 300x coverage of the tumor and the normal. Six different teams carried out mutation calling using their pipelines (different combinations of aligners, mutation callers and filters). A consensus set was generated accepting all calls made by more than three submitters. All calls made by three or fewer submitters were reviewed manually. IGV screenshots of the mutation positions, juxtaposing the normal and the tumor were generated automatically using a script developed at the DKFZ (Ivo B. please describe with some detail). The images were shared as an image library on Google+ and scored by eight reviewers manually by visual inspection. For calls that did not achieve complete agreement with the reviewers, a final decision was reached by visualizing the contentious mutation with different aligners, computing the mappability of the local region, and comparing the normal and mutant allele counts using in-house software (snape-cmp-counts), as was done above for BM1.1. The only major differences in procedure were 1) that high depth across a candidate position was also criteria for rejection and assignment to class 5, 2) multiple pipelines were used to call mutations, and 3) the candidate mutations were subjected to more manual scrutiny by the analysis team. As for BM1.1, all calls were classified according to the following criteria (Table 1): Class 1) A somatic call that is supported by >10% of the reads in the 300x set, Class 2) a somatic call that is supported by 5%-10% of the reads in the 300x set, Class 3) a somatic call that is supported by <5% of the reads in the 300x set. The estimated cut-off is 2% from where on we deemed a call unreliable due to noise. We also classified if a call fell into a region with issues of alignment (Class 4) and issues of higher than expected or lower than expected coverage for both of the sequences for the tumor and the normal (Class 5). The Gold call set was made available to all participants to review why a somatic mutation was wrongly called or missed in their respective submission.

### Evaluation of Submissions

Submissions of both CLL and MB SMMs and SIMs were evaluated against their respective Gold Sets, whose derivation is described above. For calculation of recall, the curated mutations were classified into 3 tiers according to alternate (mutation) allele frequency. Only positions were considered, not the genotypes reported. For indels, no stratification into tiers was done. For calculation of precision, all Tier 3 mutations plus ambiguously aligned mutations (class 4) were included so as to not penalize difficult to align but otherwise convincing differences between tumor and normal samples. To compare overall performance, we used a balanced measure of accuracy, the F1 score, defined as 2*(Precision*Recall)/(Precision + Recall).

Overlap calculations for the purpose of clustering and heat map generation were done using tier 3 for SSMs and tier 1 for SIMs. The Jaccard index is defined as the intersection divided by the union.

Tandem repeats were annotated with two programs. Tandem repeat finder (trf) was run on 201bp windows around each SSM and SIM call. Any repeats greater than or equal to six repeat units overlapping or immediately adjacent to the mutation position was annotated accordingly. SGA was also used to annotate homopolymers specifically, giving a richer annotation of the repeat context and change induced by the mutation.

Correspondence analysis was performed using the ‘ca’ package in R on a table where each corresponds to a genomic position at which at least one submission calls a somatic mutation in the Gold Set. The columns comprise the presence or absence in each call set, and Boolean values indicating whether certain genomic features such as repeats or presence in a blacklisted region apply, as well as sequence data such as allele frequency or depth.

Rainfall plots represent the distance for each SSM call from its immediately prior SSM on the reference genome. For each submission, the SSM set used was made of SSM called classified as True Positive or False Positive and SSM from the Gold Set Tier 4 that were absent from the submission, classified as False Negative positions.

Feature analysis was done as follows. Tandem repeats were annotated with two programs. Tandem repeat finder (trf) was run on 201bp windows around each SSM and SIM call. Any repeats greater than or equal to six repeat units overlapping or immediately adjacnt to the mutation position was annotated accordingly. SGA was also used to annotate homopolymers specifically, giving a richer annotation of the repeat context and change induced by the mutation. Mappability was calculated using the GEM package with one mismatch at both 100mer and 150mer lengths. The average 100mer and 150mer mappabilities for each mutation was calculated for a window of −90 to −10 or −140 to −10 with respect to its position, respectively. Mult100 and mult150 are defined as 1 – mappability 100 and 150. SameAF is defined as 1–2*SCOREsnape–cmp–counts–0.5)).

